# Nasopharyngeal colonisation by *Streptococcus pneumoniae* enhances host anti-viral responses to respiratory virus infection

**DOI:** 10.64898/2025.12.10.693560

**Authors:** Hanwen Hou, Julie McAuley, Ana Maluenda, Kim Mulholland, Simon Phipps, Odilia Wijburg, Catherine Satzke, Sam Manna

## Abstract

Viral-bacterial interactions during co-infection are often synergistic and can increase disease severity. However, emerging evidence indicates that some bacteria can antagonise viral infection, although the host responses driving this process remains unclear. Using infant mice co-infected with the nasopharyngeal inhabitant and pathogen *Streptococcus pneumoniae* and pneumonia virus of mice (PVM) to model antagonistic interactions, we found that prior bacterial colonisation enhances and prolongs anti-viral immune responses during co-infection, compared with viral infection alone. Transcriptomic, immunological, and histological analyses showed that pneumococcal colonisation prior to PVM infection enhanced and prolonged interferon signalling, increased anti-viral cytokine and chemokine protein levels and CD8+ cell responses. Notably, over 50% of differentially expressed host genes during co-infection were not differentially expressed in either infection alone. Our work shows that bacterial colonisation can modulate host immunity, shaping how the immune system responds to incoming viral infections, which has the potential to open novel therapeutic applications.

**Highlights:**

- Pneumococcal mono-infection in infant mice resulted in a delayed host transcriptomic response that was not detectable until 12 days post-infection.
- Host transcriptomic and immune responses to PVM were minimal except at the peak of viral replication and rapidly returned to baseline levels.
- Prior pneumococcal colonisation enhanced anti-viral immune responses to PVM infection, with more than half of the differentially expressed genes unique to co-infection.
- Pneumococcal nasopharyngeal colonisation shapes the immune response, changing how the host responds to incoming viral infections with multiple anti-viral responses that were only transiently activated during PVM mono-infection being active for a longer duration during co-infection

## Introduction

*Streptococcus pneumoniae* (the pneumococcus) is a common inhabitant of the respiratory microbiome of children, where it colonises the nasopharynx (carriage). Although this colonisation event is typically asymptomatic, it is critical for disease including otitis media, pneumonia and meningitis.^1–3^ Within the nasopharynx, pneumococci and other respiratory microbiota interact with pathogenic respiratory viruses, such as influenza and respiratory syncytial virus (RSV).^4–10^ RSV can cause severe lower respiratory tract infections and is a major cause of morbidity and mortality, placing a significant global burden on the community, healthcare system and economy.^11–13^

Pneumococcal co-infection with viruses like influenza and RSV can increase disease severity, morbidity and mortality.^14,15^ Preceding viral infection can render the host susceptible to secondary pneumococcal infection, increasing disease severity.^16,17^ Viral infection can increase pneumococcal nasopharyngeal density, adherence, dissemination and transmission in human and/or animal models.^18–23^ Multiple mechanisms contribute to these synergistic interactions, including direct pathogen-pathogen interactions or indirect modulation of host physiology.^14,24–26^ Direct binding between RSV or influenza and pneumococci can promote pneumococcal adherence to human epithelial cells and enhance pneumococcal virulence in mice.^6,27–29^ Viral infection can impair pneumococcal clearance indirectly by affecting host immune responses and promoting tissue damage.^5,6,24,27,28,30–32^ Although less studied, pneumococci can also exert antagonistic effects on viral infections. We and others have found that prior established pneumococcal colonisation decreases influenza titres, impairs replication, reduces shedding and acquisition of influenza in animal models.^10,33–35^ In a human challenge model, pneumococcal colonisation increased type I interferon (IFN) responses to a live attenuated influenza vaccine.^36^ We previously showed that pneumococcal colonisation promotes the clearance of pneumonia virus of mice (PVM, a murine analogue of respiratory syncytial virus RSV) during secondary viral infection.^35^ Interestingly, other nasal microbiota can also exert antagonistic effects on viruses.^37–43^ Despite the growing evidence on the antagonistic effects of pneumococci and other bacteria on respiratory viral infections,^10,33,36^ little is known about the host responses that may drive this process. Pneumococcal colonisation is likely to precede viral infections, particularly in low- and middle-income countries where carriage is established early in life and prevalence can reach up to 90%.^44–46^ Even in high-income countries like Australia, pneumococcal carriage can be detected before RSV infection in some infants.^47^ While most co-infection research has focused on secondary pneumococcal infection following a primary viral infection due to the clinical relevance and enhanced disease severity, little is known about how primary colonisation of commensal bacteria affects viral infections.

Here, we explored how prior bacterial colonisation shapes immune responses to subsequent viral infection. Using an infant mouse model of pneumococcal-PVM co-infection we profiled the dynamic host responses to pneumococcal and PVM mono-infection and co-infection using transcriptomics, cytokine/chemokine panels, and histological analysis. We demonstrate that pneumococcal colonisation prior to viral infection increases both the magnitude and duration of the antiviral immune responses during co-infection. Our findings provide valuable insights into potential strategies for the prevention or treatment of viral infections and highlight the potential role of nasopharyngeal microbiota in combating viral infections.

## Results

To study bacterial-mediated antagonism of respiratory viruses, we used an infant mouse model of asymptomatic pneumococcal-PVM co-infection, previously established in our laboratory, in which both pathogens are restricted to the upper respiratory tract and pneumococcal colonisation accelerates the clearance of PVM.^35^ Briefly, infant mice were inoculated intranasally with pneumococci at 5 days old, followed by PVM at 9 days old (Figure 1A). Bulk RNA-seq was conducted on nasopharyngeal tissues collected from pneumococcal mono-infected, PVM mono-infected, co-infected, and mock-infected (PBS vehicle control) mice at 13, 17, and 21 days of age, representing the early, peak, and resolution phases of PVM infection,^35^ respectively. To examine host transcriptomic responses to each infection condition in the nasopharynx, we compared the local host transcriptomic response to pneumococcal colonisation, PVM infection and co-infection with mock-infection.

**Figure 1.**
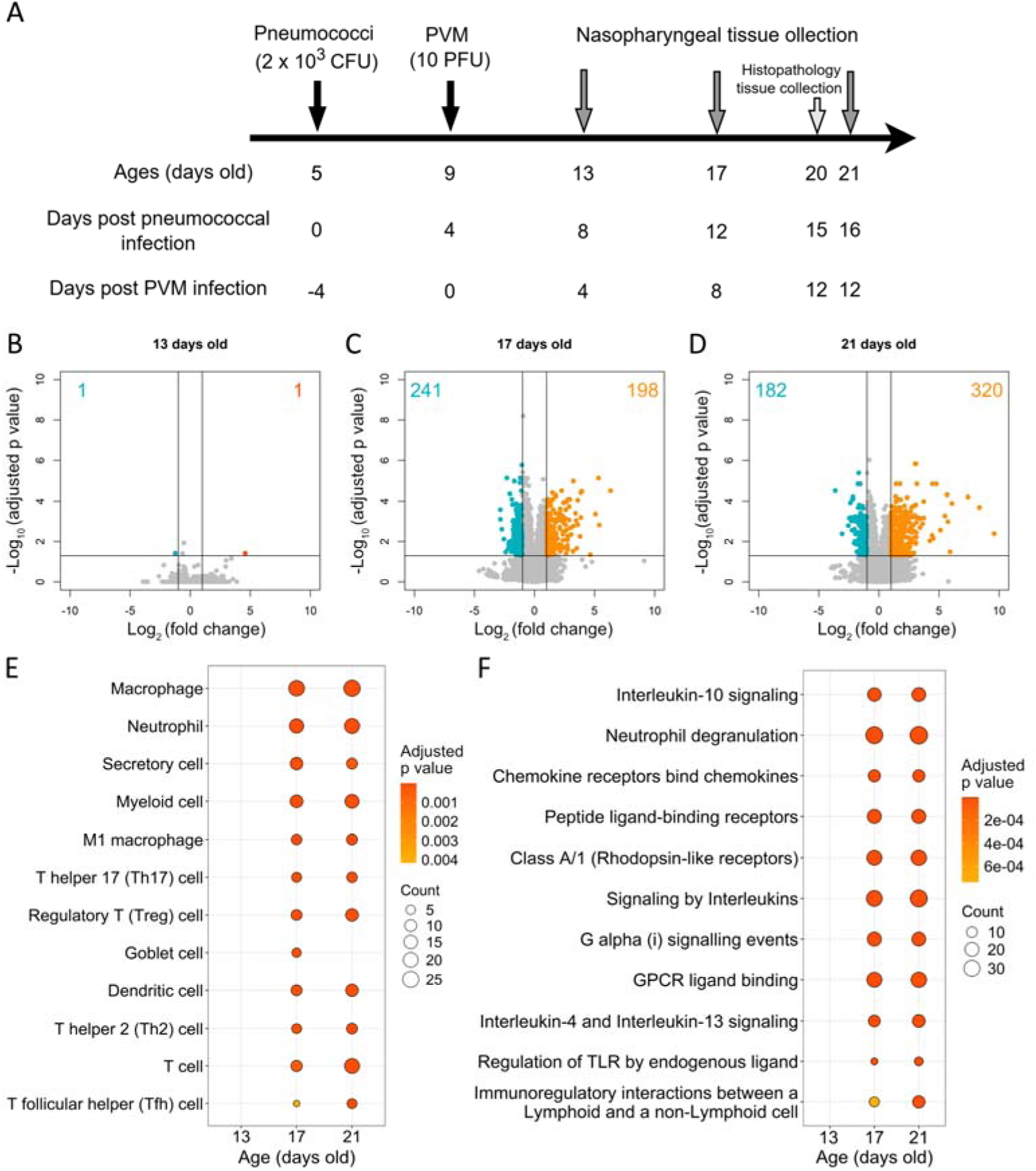
Host transcriptomic changes to asymptomatic pneumococcal colonisation. (A) Schematic representation of pneumococcal-PVM infant mouse model. Pneumococcal strain EF3030 (2 x 10^3^ CFU in 3 µl PBS) or 3 µl PBS (vehicle control) was given to mice at 5 days old intranasally without anaesthesia, followed by PVM strain J3666 (10 PFU in 3 µl PBS) or 3 µl PBS at 9 days old. Mice were euthanized and nasopharyngeal tissues were collected at 13, 17 and 21 days old. Mouse heads were collected at 20 days old for downstream histopathology examination. (B-D) Volcano plots of differentially expressed genes (DEGs) during pneumococcal mono-infection at 13 (B), 17 (C) and 21 (D) days old compared with mock-infected mice. X and Y axes represent log_2_ (fold change) (LFC) and -log_10_ (adjusted p value). Genes with LFC <-1 or LFC >1 with adjusted p value < 0.05 are considered DEGs. Blue and orange indicate down-regulated and up-regulated genes, respectively. (E and F) Over-representation analysis of pneumococcal mono-infection for most enriched cell types (E) and pathways (F) at 13, 17 and 21 days old compared with mock-infected mice. The count indicates the number of DEGs identified within the gene set corresponding to each cell type or pathway in the reference database.

### Host transcriptomic changes to asymptomatic pneumococcal colonisation

At 13 days of age (8 days after pneumococcal infection), only two host genes were differentially expressed, with one upregulated and the other downregulated (Figure 1B). In contrast, 439 (198 up, 241 down) and 502 (320 up, 182 down) host genes were differentially expressed at 17 and 21 days old, respectively, compared with mock-infected controls (Figure 1C-D, Table S1). Host genes upregulated in response to pneumococcal colonisation included those involved in inflammation and bacterial defence, such as *Il23* and *Tnf*, with the host transcriptome showing similar patterns at 17 and 21 days old (Figure S1A, Table S1).

To identify cell types and biological pathways enriched in response to pneumococcal colonisation alone, an over-representation analysis was performed by grouping differentially expressed genes (DEGs) according to their biological pathways in the Reactome database.^48^ Among the DEGs identified, genes associated with innate immune cell types, including neutrophils, macrophages and dendritic cells (DCs), were predicted to be enriched at both 17 and 21 days old (Figure 1E). Similarly, pathway prediction based on DEGs suggested an enrichment of innate immune response-related pathways, such as neutrophil degranulation (Figure 1F). A range of genes related to neutrophil recruitment and trafficking were significantly enriched, including *Cxcl1*, *Cxcl2*, *Cxcr2*, *Ccl3*, *Ccl4* and *Csf3,* as well as anti-microbial functions such as *Elane* (neutrophil elastase), *Ctsg* (cathepsin G) and *Prtn3* (proteinase 3). *Cxcl2* was the only DEG enriched at all time points tested; CXCL2 mediates neutrophil adhesion and attracts neutrophils to the site of bacterial infection to promote host defence.^49–51^

In addition to innate cells, adaptive immune cells were also predicted to be enriched during pneumococcal mono-infection, specifically T cells, including T helper 17 (Th17) cells and regulatory T (Treg) cells (Figure 1E). The signalling pathway of IL-10, a potent anti-inflammatory cytokine mainly produced by CD4+ T cells, including Treg cells,^52^ was enriched during pneumococcal colonisation (Figure 1F). In addition, we also observed an enrichment of Th2-related genes and pathways, including *Ccr4/Ccl22* and genes related to IL-4 and IL-13 signalling pathways (Figure 1F), indicative of a Th2 response.^53^ Of note, during pneumococcal mono-infection, only one immunoglobulin gene was up-regulated at 17 days old, while 31 immunoglobulin genes were up-regulated at 21 days old (Table S1). Four genes were related to IgG response, and one was associated with IgA response, suggesting B cell and antibody responses at 21 days old (Table S1). Interestingly, pneumococcal colonisation also induced differential expression of genes associated with natural killer (NK) cells and CD8+ T cells at both 17 and 21 days old (Table S1). Although these two cell types could contribute to controlling pneumococcal infection,^54,55^ they are primarily known for their anti-viral roles.^56,57^

### Host transcriptomic changes to PVM infection

Compared with responses induced by pneumococcal colonisation, the host response to PVM mono-infection was relatively muted. At 13 days old and 21 days old, no genes were differentially expressed. At 17 days old (the peak of PVM load),^35^ 222 genes (166 up, 56 down) were differentially expressed during PVM mono-infection (Figures 2A-C and Table S2). PVM mono-infection induced a distinct expression profile compared with mock infection, characterised by the up-regulation of genes associated with inflammation and anti-viral responses, such as *Isg15*, *Cxcl10*, and *Ifng* (Figure S1B).

**Figure 2.**
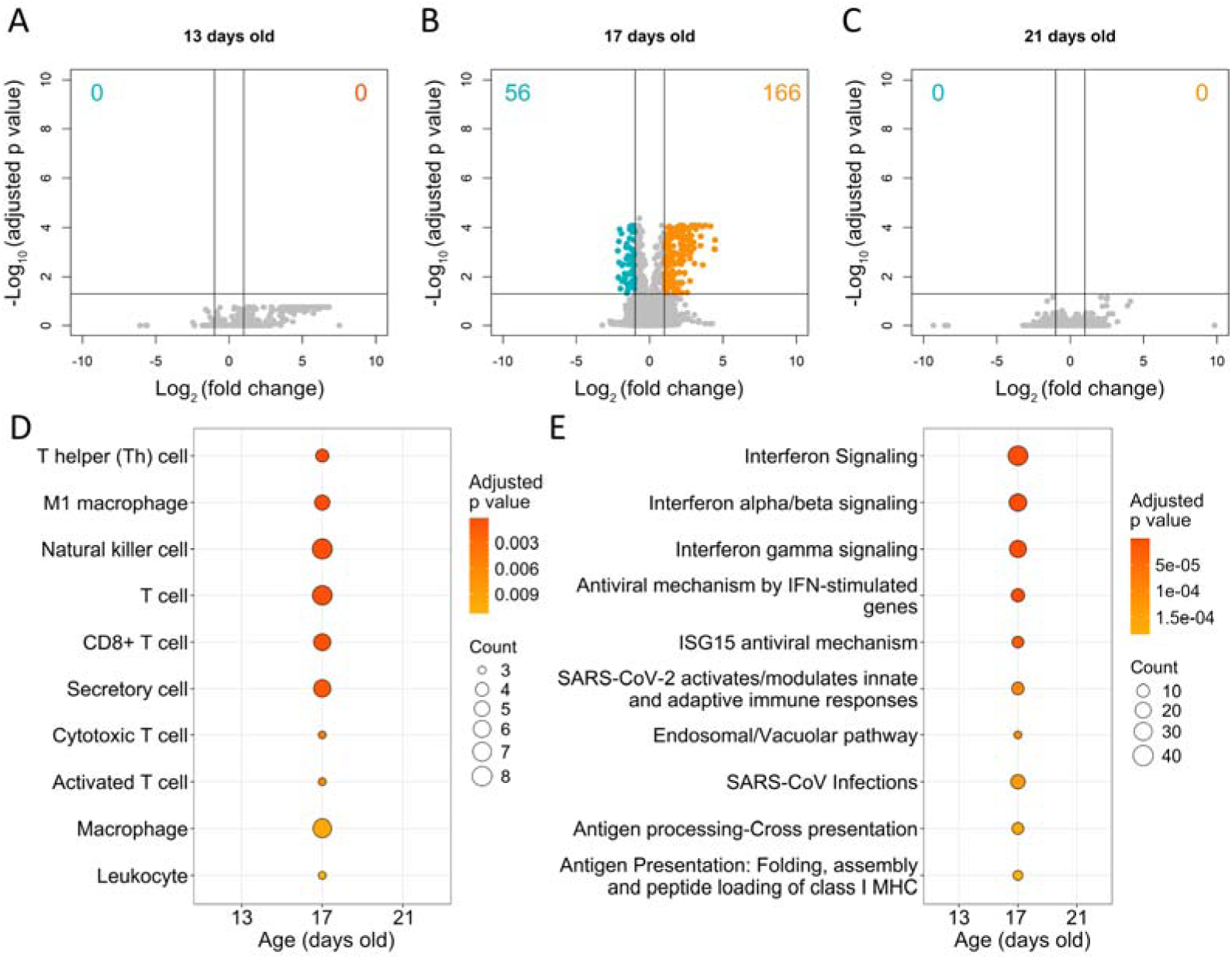
Host transcriptomic changes to PVM infection. (A-C) Volcano plot of DEGs during PVM mono-infection at 13 (A), 17 (B) and 21 (C) days old compared with mock-infected mice. X and Y axes represent LFC and -log_10_ (adjusted p value). Genes with LFC <-1 or LFC >1 with adjusted p value < 0.05 are considered DEGs. Blue and orange indicate down-regulated and up-regulated genes, respectively. (D and E) Over-representation analysis of PVM mono-infection for most enriched cell types (E) and pathways (F) at 13, 17 and 21 days old compared with mock-infected mice. The count indicates the number of DEGs identified within the gene set corresponding to each cell type or pathway in the reference database.

At 17 days old, cell type prediction suggested an enrichment of both innate immune cells, which includes neutrophils, macrophages, DCs and NK cells, and adaptive immune cells consisting of both CD4+ and CD8+ T cells (Figure 2D). Genes encoding effector molecules (including granzymes and perforin) of CD8+ T cells and NK cells were enriched, two key anti-viral cell types that play a crucial role in defending against RSV and influenza infection in neonates.^58–63^ IFN (IFN-α/β and IFN-γ) pathways were enriched during PVM mono-infection (Figure 2E), which are critical parts of anti-viral immune defence by activating downstream signalling and multiple anti-viral mechanisms.^64^ The expression of IFNs and TNF-α-related anti-viral genes and pathways was elevated at 17 days old, including guanylate-binding proteins *Gbp* genes, 2’,5’-oligoadenylate synthetase *Oas* genes, *Isg15*, *Mx1/Mx2* and *Cxcl9/Cxcl10/Cxcl11*.

### Pneumococcal colonisation promotes anti-viral transcriptomic responses during PVM infection

Compared with mock infection, a total of 0, 989 (659 up, 330 down) and 1350 (482 up, 868 down) genes were differentially expressed at 13, 17 and 21 days old during co-infection, respectively (Figure 3 A-C and Table S3). This number of DEGs was higher than with either pneumococcal or PVM mono-infection at both 17 and 21 days old (Figure 1B-D, Figure 2A-C). The transcriptomic response during co-infection partially overlapped with either mono-infection, with a higher proportion of DEGs in common with pneumococcal colonisation (Figure 3D-F and Table S3). During co-infection, of the 989 DEGs found at 17 days old, 318 (32.2%) and 195 (19.7%) genes were in common with pneumococcal or PVM mono-infection, respectively (Figure 3E). Similarly, at 21 days old, among 1350 DEGs, 439 (32.5%) DEGs were found in both pneumococcal mono-infection and co-infection, while 0 DEGs were in common between PVM mono-infection and co-infection (Figure 3F).

**Figure 3.**
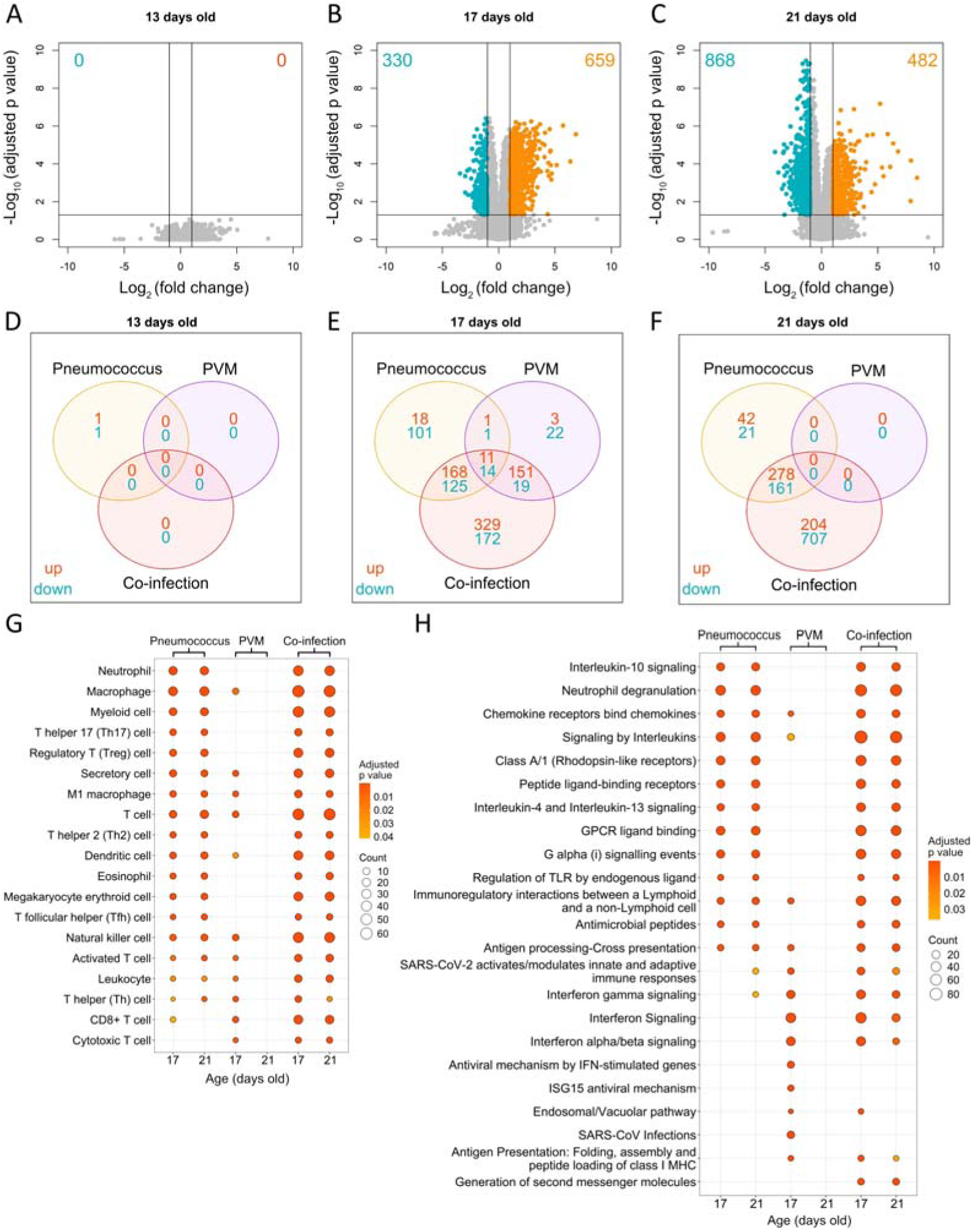
Host transcriptomic changes to co-infection. (A-C) Volcano plot of DEGs during co-infection at 13 (A), 17 (B) and 21 (C) days old compared with mock-infected. X and Y axes represent LFC and -log10 (adjusted p value). Genes with LFC <-1 or LFC >1 with adjusted p value < 0.05 are considered DEGs. Cyan, grey and orange indicate down-regulated, not significant and up-regulated genes, respectively. (D-F) Venn diagram of overlapping and non-overlapping genes from co and mono-infection at 13 (D), 17 (E) and 21 (F) days old. (G and H) Over-representation analysis of co-infection and mono-infections for most enriched cell types (G) and pathways (H) at 17 and 21 days old compared with mock-infected mice. Data at 13 days old was not plotted due to no over-represented cell types or pathways. The count indicates the number of DEGs identified within the gene set corresponding to each cell type or pathway in the reference database.

To identify host responses contributing to pneumococcal-mediated PVM clearance, we analysed DEGs unique to co-infection compared with mock controls. As expected, compared to mock controls, co-infection caused an enhanced breadth of gene expression at heightened levels. Particularly, at 17 days old, 501 (50.7%) of DEGs were unique to co-infection, including *Csf2*, *Ccl2*, and *Cxcl13* (Figure 3E, Figure S2A). At 21 days old, 911 DEGs (67.5%) were unique to co-infection, with no overlap between PVM mono-infection and co-infection (Figure S2B). A range of DEGs unique to co-infection at 21 days old, including *Gpb* and *Oas* genes, were detected earlier during PVM mono-infection at 17 days old.

Over-representation analysis of DEGs during co-infection identified enriched cell types and pathways, most of which overlapped with those observed in mono-infections (Figure 3G and 3H). During co-infection, DEGs associated with innate immune cell populations, including neutrophils, macrophages, dendritic cells and NK cells, were enriched compared with mock controls at both 17 and 21 days of age (Figure 3G). Innate immune functions, such as neutrophil degranulation and antigen presentation, were also enriched during co-infection (Figure 3H). Pathways related to adaptive immune cells were enriched, including the CD8+ T cell, which is one of the main contributors to the clearance of PVM and RSV.^57,60,65^ Both anti-bacterial and anti-viral responses were seen during co-infection (Figure 3H). Anti-viral responses, including IFN signalling and antigen presentation pathways, persisted at 21 days during co-infection but not during PVM mono-infection, indicating that pneumococcal colonisation may prolong these anti-viral responses. In addition, over-representation analysis based on DEGs that were unique to co-infection indicated the enrichment of various anti-viral cell types and pathways, including CD8+ T cells and NK cells (Figure S2A), and IFN-γ signalling pathways at both 17 and 21 days old (Figure S2B).

### Pneumococcal colonisation promotes anti-viral responses during co-infection

Our data suggest pneumococcal colonisation triggers changes in the host transcriptome, including upregulation and persistence of anti-viral pathways during co-infection. To validate the RNA-seq results at the protein level, concentrations of selected cytokines and chemokines were measured at 13, 17 and 21 days old during mock, pneumococcal mono-infection, PVM mono-infection or co-infection (Figure 4 and Figure S3). Cytokines involved in neutrophil chemotaxis, survival, proliferation, and function, such as CXCL1 (KC), CXCL2 (MIP-2α), and CSF3 (G-CSF), were elevated during both pneumococcal mono-infection and co-infection at all time points (Figure 4A-C), whereas at 13 days old, only CXCL2 showed transcriptomic changes during pneumococcal mono-infection (Table S1). The levels of CCL5 (RANTES), CCL2 (MCP-1), CCL3 (MIP-1α), and CCL4 (MIP-1β), which affect monocyte/macrophage function,^66–69^ were higher during co-infection at 17 days old compared with PVM mono-infection (Figure 4D-G). CCL5 increased to a similar level for PVM mono-infection and co-infection at 21 days old, while CCL2, CCL3 and CCL4 remained higher during co-infection at both 17 and 21 days old. The level of IL-6, which could affect both innate and adaptive immune responses, recruit macrophages and affect T cell subset commitment,^70^ was highest at 17 days old during co-infection (Figure 4H).

**Figure 4.**
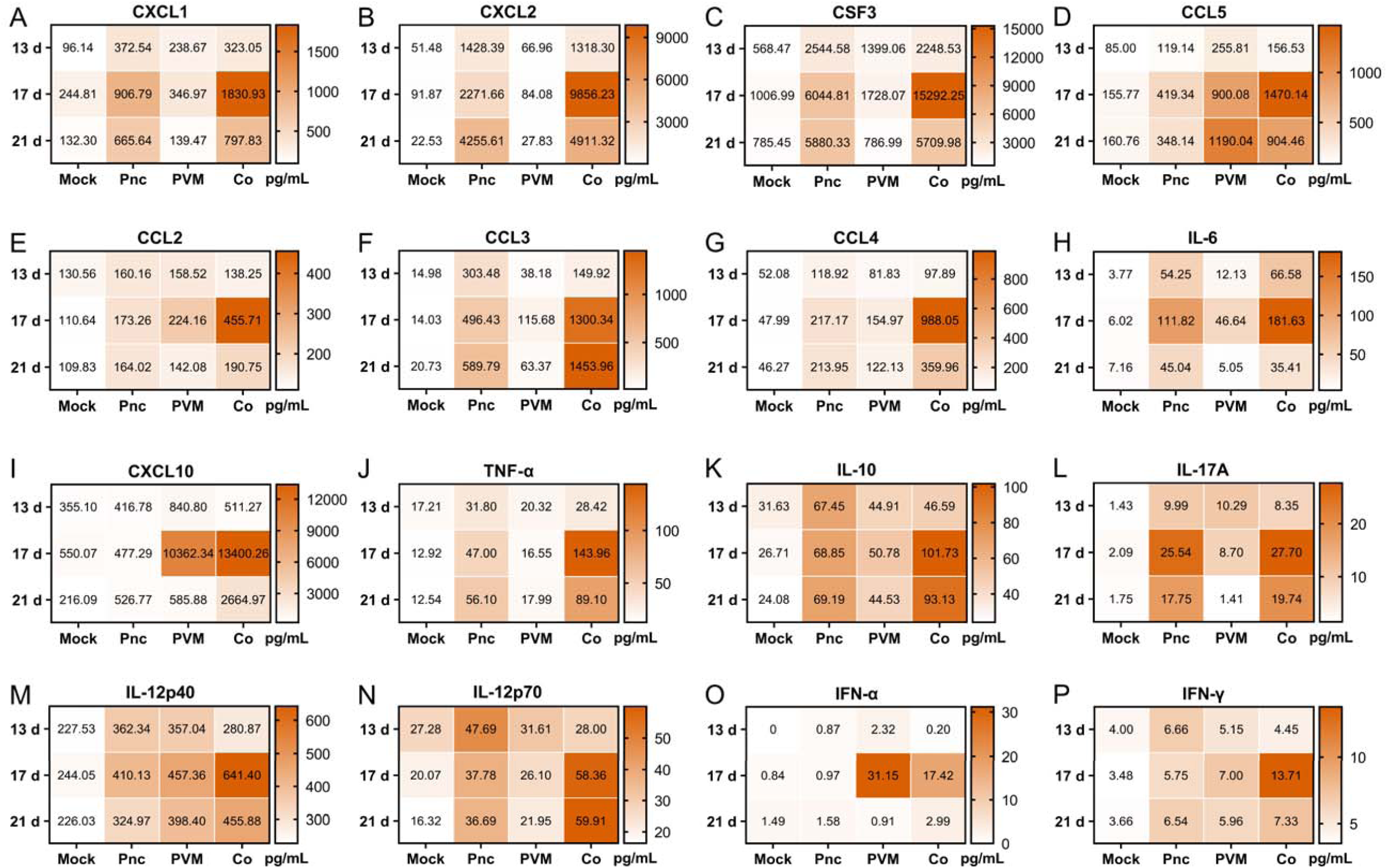
Cytokine levels during pneumococcal-PVM co-infection over time. (A to P) Concentrations of critical cytokines and chemokines involved in mono-infections and co-infections at each time point. Concentrations of cytokines and chemokines in nasopharyngeal homogenates of 13, 17 or 21-day-old mice that were mock infected (Mock), pneumococcal mono-infected (Pnc), PVM mono-infected (PVM), or co-infected (Co). The colour of each block corresponds to the colour bar and represents the mean concentration of six samples per condition per time point.

Pro-inflammatory cytokines that contribute to anti-viral responses and T cell responses had elevated concentrations during co-infection, including CXCL10 (IP-10), TNF-α, IL-1α and IL-1β (Figure 4I-J, Figure S3A-B). Consistent with the host transcriptome, CXCL10 remained at a high concentration during co-infection at 17 days old and dropped to a lower level at 21 days old, while it was higher during co-infection at 21 days old compared with PVM mono-infection (Figure 4I). TNF-α showed a similar trend as neutrophil-related cytokines and chemokines, with its concentration being higher during pneumococcal mono-infection and co-infection compared with PVM mono-infection or mock at both 17 and 21 days old, with the highest seen during co-infection at 17 days old (Figure 4J).

The level of IL-10 was higher during co-infection compared with mono-infection at 17 and 21 days old (Figure 4K). IL-17A, a pro-inflammatory cytokine involved in pneumococcal clearance,^71,72^ was elevated during both mono- and co-infections, with higher levels observed during pneumococcal mono-infection and co-infection (Figure 4L). Concentrations of IL-12p40 and IL-12p70 were slightly elevated during co-infection compared with either mono-infections or mock at 17 and 21 days old (Figure 4M and 4N). IFN-α, a key anti-viral cytokine, peaked at 17 days old during both PVM mono-infection and co-infection, with higher levels observed in PVM mono-infection (Figure 4O). The levels of IFN-γ remained stable at all timepoints, while being higher during co-infection compared with mono-infection at 17 days old (Figure 4P). Of note, consistent with the Th2-biased neonatal immune system,^73–76^ a range of Th2/eosinophil-related cytokines and chemokines was observed in all conditions, including IL-4, IL-13, CCL11 (Eotaxin), IL-5 and IL-9 (Figure S3D-I).

### Quantification and identified of cell types by immunohistochemistry

Next, to examine cells recruited to the nasopharynx during each condition, we performed immunohistochemistry staining on sections of mice nasopharynx under each infection condition at 20 days old, at which point the pneumococcal-mediated antagonism on PVM occurs. Samples were stained for LyG6, CD4 and CD8A for quantification and imaging. LyG6, CD4 and CD8A are mainly expressed by neutrophils, CD4+ and CD8+ T cells, respectively.^77–79^ Consistent with transcriptomic and cytokine findings, LyG6+ cells were abundant during pneumococcal mono-infection and co-infection, comprising ∼5% of all nasopharyngeal cells, but accounted for <1% of cells during PVM mono-infection and mock infection (Figure 5A). Ly6G+ cells were distributed throughout the mucosal and submucosal layers and formed aggregates within the exudate in the nasal cavity during pneumococcal mono-infection and co-infection, whereas they appeared only sporadically during mock and PVM mono-infection (Figure 5B-E). CD4+ cells were rarely observed in mock-infected mice (<0.5%) but increased in number in all infection conditions, with higher numbers in PVM mono-infected (∼1%), pneumococcal mono-infected (∼2%), and co-infected (∼1.5%) mice, appearing occasionally throughout the tissue (Figure 6). Similarly, CD8+ cells were infrequent in mock-infected mice (<0.2%) but were more abundant during infection, with levels of ∼0.6% in PVM mono-infection, ∼0.5% in pneumococcal mono-infection, and approximately ∼1% during co-infection, frequently found among the mucosal epithelial layer (Figure 7).

**Figure 5.**
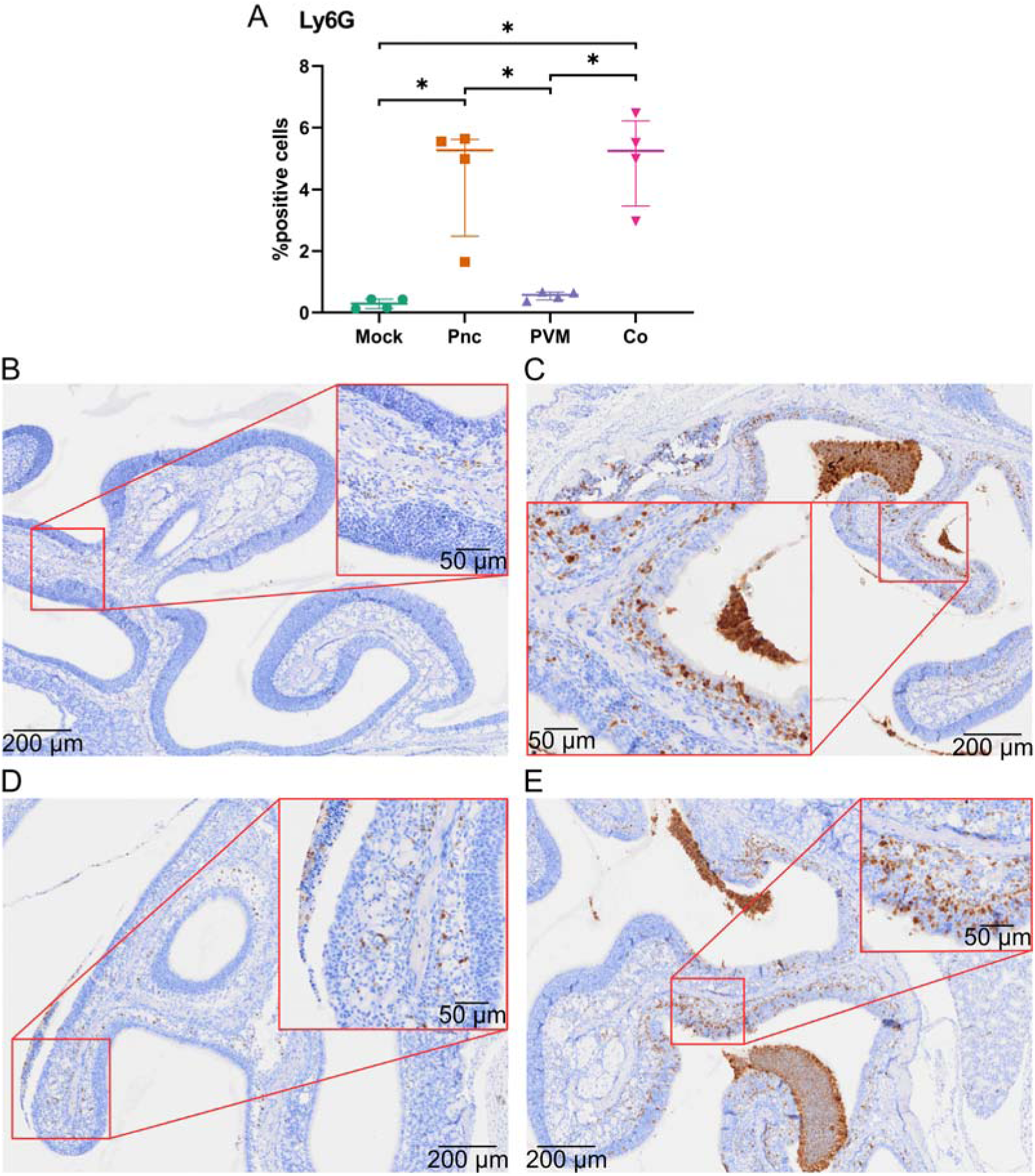
Count and percentage of Ly6G-positive cells identified by immunohistochemistry. (A) percentage of Ly6G-positive cells out of all cells from mice that were mock infected (Mock), pneumococcal mono-infected (Pnc), PVM mono-infected (PVM), or co-infected (Co). Data presented as median ± interquartile range (IQR). The p values for carriage density comparisons were calculated using the Mann-Whitney test, * represents p < 0.05. (B-E) representative images of mice nasopharynx during mock (B), pneumococcal mono-infection (C), PVM mono-infection (D) and co-infection (E) at 20 days old. Paraffin blocks containing decalcified murine nasal cavities were submitted for sectioning. Sections were prepared at 4 μm, and all sections were stained with DAB chromogenic immunohistochemistry. Cell counts were performed in the area surrounding the hard tissue support beneath the mucosa and submucosa. Intranasal cavity spaces, including mucus, cellular aggregates, and exudates, were excluded from the analysis. After staining, sections were scanned using a VS200 scanner, under brightfield mode, with a 20x objective. Scale bar_=_200 μm, insets (denoted by boxes) are at 50 μm.

**Figure 6.**
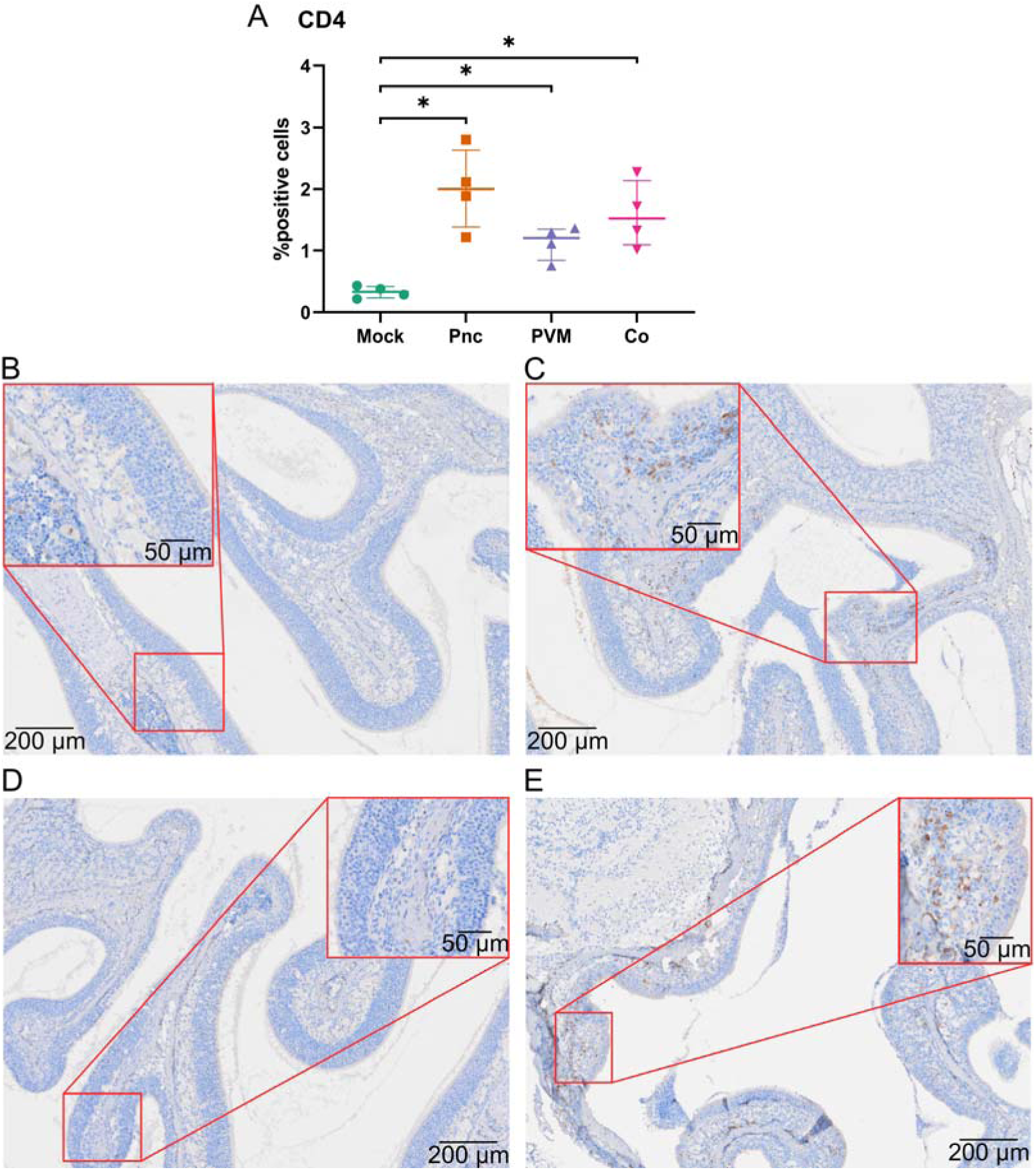
Count and percentage of CD4-positive cells identified by immunohistochemistry. (A) percentage of CD4-positive cells out of all cells from mice that were mock infected (Mock), pneumococcal mono-infected (Pnc), PVM mono-infected (PVM), or co-infected (Co). Data presented as median ± interquartile range (IQR). The p values for carriage density comparisons were calculated using the Mann-Whitney test, * represents p < 0.05. (B-E) representative microscopic image of mice nasopharynx during mock (B), pneumococcal mono-infection (C), PVM mono-infection (D) and co-infection (E) at 20 days old. Paraffin blocks containing decalcified murine nasal cavities were submitted for sectioning. Sections were prepared at 4 μm, and all sections were stained with DAB chromogenic immunohistochemistry. Cell counts were performed in the area surrounding the hard tissue support beneath the mucosa and submucosa. Intranasal cavity spaces, including mucus, cellular aggregates, and exudates, were excluded from the analysis. After staining, sections were scanned using a VS200 scanner, under brightfield mode, with a 20x objective. Scale bar_=_200 μm, insets (denoted by boxes) are at 50 μm.

**Figure 7.**
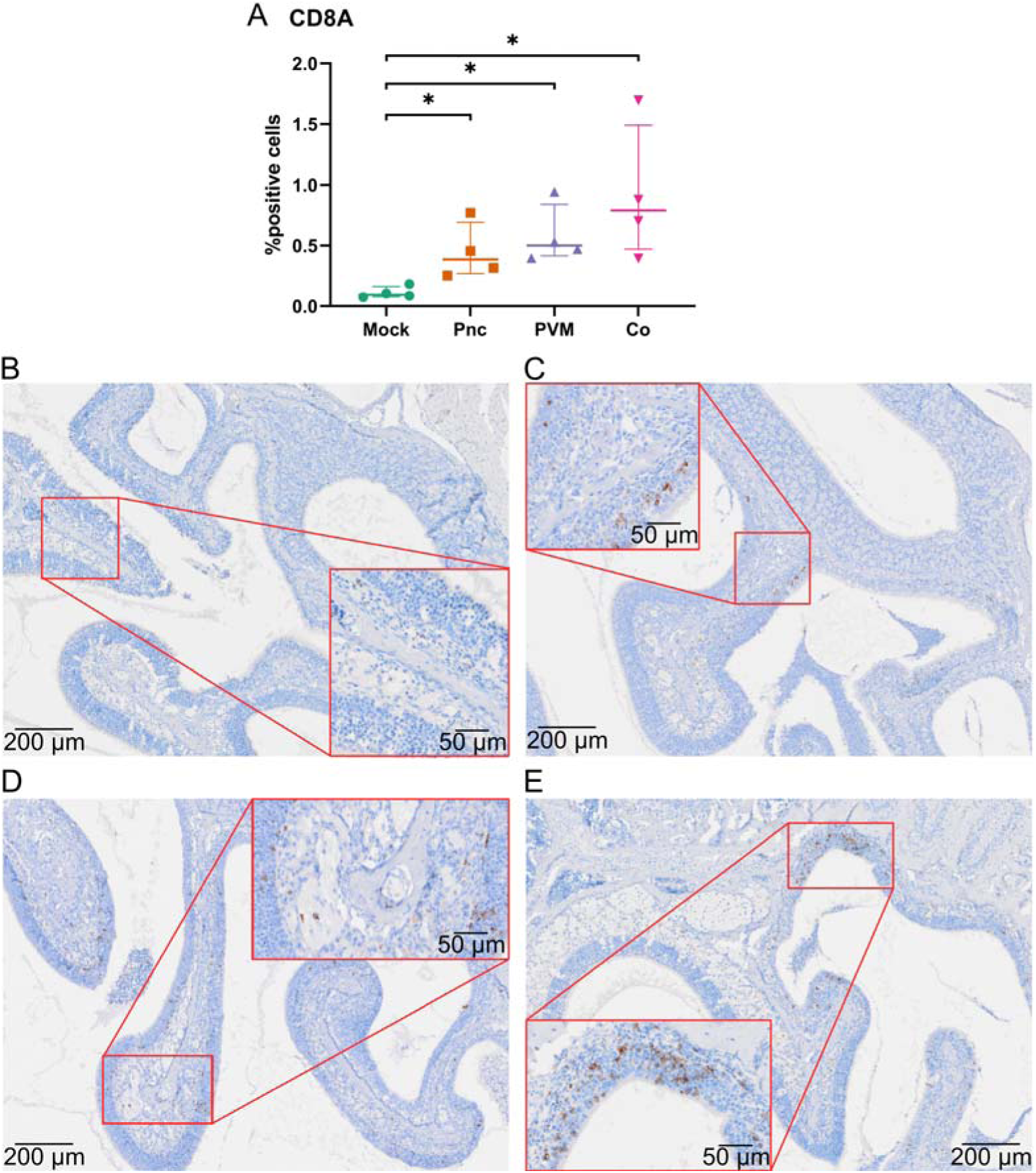
Count and percentage of CD8A-positive cells identified by immunohistochemistry. (A) percentage of CD8-positive cells out of all cells from mice that were mock infected (Mock), pneumococcal mono-infected (Pnc), PVM mono-infected (PVM), or co-infected (Co). Data presented as median ± interquartile range (IQR). The p values for carriage density comparisons were calculated using the Mann-Whitney test, * represents p < 0.05. (B-E) representative microscopic image of mice nasopharynx during mock (B), pneumococcal mono-infection (C), PVM mono-infection (D) and co-infection (E) at 20 days old. Paraffin blocks containing decalcified murine nasal cavities were submitted for sectioning. Sections were prepared at 4 μm, and all sections were stained with DAB chromogenic immunohistochemistry. Cell counts were performed in the area surrounding the hard tissue support beneath the mucosa and submucosa. Intranasal cavity spaces, including mucus, cellular aggregates, and exudates, were excluded from the analysis. After staining, sections were scanned using a VS200 scanner, under brightfield mode, with a 20x objective. Scale bar_=_200 μm, insets (denoted by boxes) are at 50 μm.

## Discussion

There is increasing evidence that there can be antagonistic relationships between bacteria and viruses,^10,33,35,80^ although the host response driving these interactions remains largely unknown. In this study, we investigated the effect of pneumococci on PVM infection as a model where we have previously found an antagonistic effect of pneumococci on PVM. We found that pneumococcal colonisation induced sustained upregulation of neutrophil-associated genes and proteins, including chemoattractants, activation markers, and effector molecules. Immune responses to PVM peaked concurrently with the peak viral loads and then declined rapidly as the infection was being resolved.^35^ In contrast, co-infection induced a unique transcriptomic signature, as more than half of DEGs were not enriched in either mono-infection. During co-infection, pneumococcal colonisation enhanced both the magnitude and duration of anti-viral cytokine and chemokine responses compared with PVM mono-infection and increased the number of immune cells infiltrating in the nasopharynx.

Pneumococci commonly colonise infants and young children and the duration of carriage can exceed 200 days.^81–84^ Most studies on pneumococci and respiratory viruses use adult models of infection. However, the infant immune system differs from that of adults, being characterised by an immunosuppressive environment.^85,86^ The responses to either pneumococci or PVM are poorly characterised in infant hosts, and even less is known regarding co-infections and how infants respond in this context. To address these knowledge gaps, we performed a detailed investigation of the local transcriptomic changes to pneumococcal nasopharyngeal colonisation and PVM infection using an infant mouse model. Neutrophils are among the earliest and most critical components of the innate immune response to pneumococcal colonisation and lung infection.^49,87^ Our data suggest that CXCL2 and CCL3, which have not previously been shown to be induced during pneumococcal carriage, may contribute to the neutrophil response in infants. Interestingly, several chemokines and cytokines with known anti-viral roles, including CCL3, CCL5, and TNF-α,^88–97^ were enriched during pneumococcal mono-infection at both the RNA and protein level, potentially priming earlier anti-viral responses that could antagonise subsequent viral infections.

To characterise PVM mono-infection in the upper respiratory tract, we comprehensively profiled host responses at both RNA and protein levels. During PVM mono-infection, immune responses were the highest at 8 days post-infection and declined afterwards in the nasopharynx, which coincides with the expected peak of virus load.^35^ We observed upregulation of several key anti-viral responses, including ISG15, MX1/MX2 and IFI16.^98–102^ *Cxcl9*, *Cxcl10* and *Cxcl11* were among the most upregulated genes, and encode for chemokines responsible for attracting macrophages, monocytes, NK and T cells during viral infection.^103,104^ In our study, CXCL10 protein levels were higher during co-infection at both 17 and 21 days old. CXCL10 is predominantly induced by IFN-γ, which can promote RSV clearance.^105,106^

RSV-pneumococcal co-infection also leads to an earlier expression of CXCL10 and reduced RSV loads compared with RSV alone in human monocyte-derived macrophages.^107^ The heightened and prolonged expression of CXCL10 during co-infection may play a role in the pneumococcal-mediated antagonistic effects on PVM in our model.

Co-infection of bacterial and viral pathogens could synergistically increase immune responses, leading to higher anti-viral responses. IFNs play critical roles in innate defences against respiratory viral infections, which activate anti-viral mechanisms such as the OAS, GBP and MX proteins.^108–113^ The number of DEGs in the IFN-γ signalling pathway was higher at 17 and 21 days old during co-infection, compared with either mono-infection, characterised by the DEGs uniquely found during co-infection. A low level of IFN-γ and IFN-γ-producing cells is associated with delayed RSV clearance and increased disease severity in humans and infant mice.^114–117^ Both pneumococci and respiratory viruses, including RSV and influenza, can increase the level of IFN-γ.^118–121^ Our data showed that co-infection led to extended IFN signalling (including IFN-α/β and IFN-γ), with increased concentrations and expression of IFNs and IFN-regulated cytokines and chemokines compared with PVM mono-infection. Notably, these cytokines and chemokines include IFN-γ-inducible TNF-α, CXCL10 and CCL3, which can contribute to anti-viral responses.^93–97,122–124^ Thus, by enhancing and prolonging IFN signalling and downstream pathways, pneumococcal colonisation may promote PVM clearance during co-infection.

Over 50% of DEGs identified during co-infection at both 17 and 21 days of age were not enriched in either mono-infection. The uniquely enriched genes during co-infection were associated with NK cells, CD8+ T cells, and IFN-γ signalling pathways, suggesting that these immune responses may play key roles in pneumococcal-mediated antagonism of PVM. IFNs also promote anti-viral responses of NK and CD8+ T cells, which are two of the most critical cell types mediating RSV and PVM clearance, especially in neonates.^60,115,125,126^ NK cells are the main IFN-γ producing cells during RSV infection, and increased NK cell cytotoxicity and IFN-γ production correlate with increased numbers of CD8+ T cells in RSV vaccinated mice.^127^ NK cells peak around the same time as PVM replication peak, and it may play a similar role as CD8+ T cells to induce cell death and limit PVM replication, therefore promoting clearance.^128^ The elevated IFN-γ levels and upregulation of IFN-γ-associated DEGs during co-infection may contribute to the increased abundance of CD8+ cells observed at 20 days of age compared with mono-infection, and CD8+ T cell responses are known to promote effective control and clearance of RSV and PVM.^57,60,129–133^ In addition to IFNs, TNF-α can be secreted by activated T cells and NK cells, promoting the activation and proliferation of both CD4+ and CD8+ T cells, and contributing to anti-viral responses.^93–95^ CCL3, CCL4 and CCL5, which are produced by activated NK, CD8+ T, and CD4+ T cells, can synergise with IFN-γ to upregulate TNF-α and promote the transition of innate immunity to antigen-specific CD8+ T cell adaptive immunity.^66^ Higher levels of these cytokines and chemokines at 17 days old as a result of pneumococcal colonisation may increase CD8+ T cell responses or promote an earlier transition to antigen-specific CD8+ T cell adaptive immunity that is effective against RSV infection.

Overall, our findings suggest that pneumococcal colonisation triggers earlier and stronger innate immune responses, including a range of anti-viral cytokines, chemokines, and innate immune cells such as NK cells. These elevated innate signals may promote earlier or more robust activation of CD8+ T cells, thereby enhancing and prolonging adaptive responses, such as CD8+ T cell recruitment and activation, which could explain the accelerated PVM clearance we observed previously.^35^.

In summary, our work examined how a bacterial member of the respiratory microbiome can antagonise respiratory viral infections and demonstrated that colonising bacteria modulate the infant host response in ways that limit subsequent viral infection. Our data suggest that pneumococcal colonisation resulted in elevated and prolonged anti-viral responses, including cytokines and chemokines linked to innate and adaptive immunity, which may contribute to pneumococcal-mediated acceleration of PVM clearance. Despite advances in the prevention and treatment of respiratory viral infections, current strategies remain inadequate, emphasising the need for more effective therapeutics and preventive measures. Our work highlights that certain members of the microbiome may play a critical role in modulating anti-viral immune responses, potentially contributing to controlling incoming viral infections and thereby influencing the outcome of those infections.

## STAR Methods

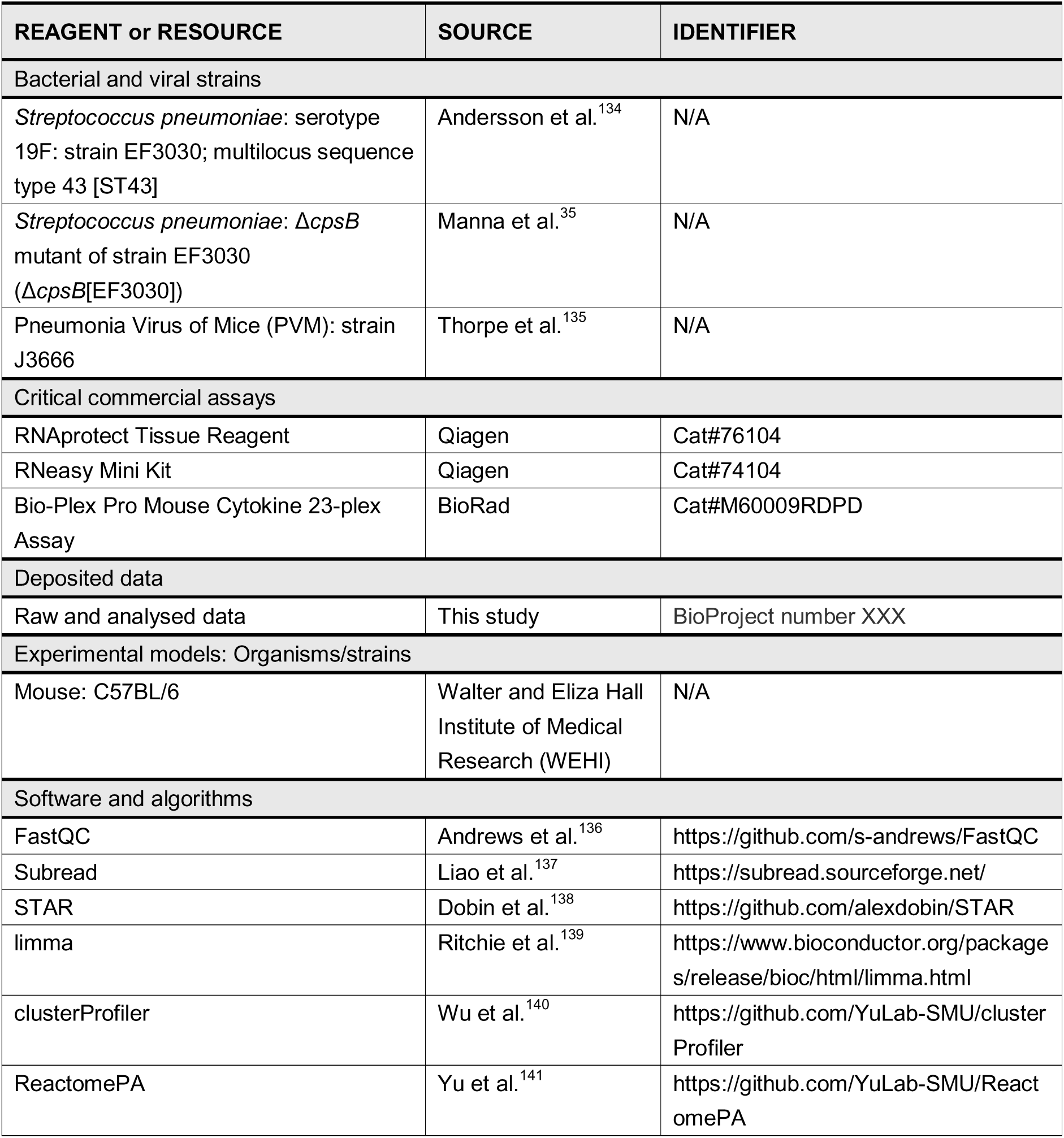
Key resources table

### Resource availability

#### Lead contact

Further information and requests for resources and reagents should be directed to and will be fulfilled by the lead contact, Catherine Satzke

#### Materials availability

This study did not generate new unique reagents.

#### Data and code availability

RNA sequences have been deposited at the European Nucleotide Archive with project accession number PRJEB104213

This paper does not report original code.

Any additional information required to reanalyse the data reported in this paper is available from the lead contact upon request.

### Experimental model and study participant details

#### Infant mice model

All mouse studies were carried out with the approval of the Murdoch Children’s Research Institute (MCRI) Animal Ethics Committee (ethics numbers A832 and A936), in compliance with the Australian code for the care and use of animals in scientific research.^142^ The infant mouse model used in this study was established previously.^35^ In brief, C57BL/6 pups were raised under SPF conditions and were given 2[×[10^3^ CFU of pneumococci (strain EF3030) in 3 μl PBS intranasally without anaesthesia at 5 days old, followed by 10 PFU of PVM (strain J3666) in 3 μl PBS at 9 days old. The mock control and mono-infection groups received 3 μl PBS as a vehicle control. At 13, 17 and 21 days old, mouse nasopharyngeal tissue was collected in 1.5 ml RNAprotect Tissue Reagent (Qiagen) or 500 µl RPMI 1640 medium and homogenised for downstream RNA extraction or cytokine analysis, respectively. Mouse heads were collected at 20 days old, which were prepared and sectioned as previously described^35^ for downstream histology analysis.

### Method details

#### Bacterial and viral strains

Pneumococcal infectious stocks were prepared as previously described.^35^ In brief, pneumococci (strain EF3030) were grown in THY broth at 37°C with 5% CO_2_. After pneumococci reached ∼0.4 OD_600_, glycerol was added to reach 8% (vol/vol) final concentration for storage at −80°C. PVM (strain J3666) was purified from mouse lung homogenates as described previously.^143^

#### RNA extraction and sequencing

Host RNA was extracted from mice nasopharyngeal tissues of each condition at 13, 17, and 21 days old using the RNeasy Mini Kit (Qiagen) as per the manufacturer’s instructions. The quality of extracted host RNA was checked with the Agilent TapeStation system; RNA with an RNA integrity number (RIN) score over 8 was used in this study. Library preparation was performed using the Illumina TruSeq stranded mRNA kit and sequenced on the Illumina NovaSeq 6000 sequencer (2×150bp) at 30M reads.

#### RNA-seq analysis

Sequencing quality control was performed using FastQC^136^ V0.11.8, followed by alignment to the mouse reference genome assembly GRCm39 (RefSeq assembly GCF_000001635.27) using STAR^138^ aligner V2.7.10a and counting using featureCounts from the Subread^137^ package V2.0.0; reads with a Phred quality score less than 10 were removed during alignment. Counts were normalised and analysed using the limma^139^ package V3.56.1 in R V4.3.1.

Genes were considered differentially expressed if a gene had a log_2_ fold change > 1 or log_2_ fold change < -1 with a Benjamini-Hochberg adjusted p value < 0.05. Over-representation analysis was performed using R packages ClusterProfiler^140^ V4.10.1 and ReactomePA^141^ V1.46.0 with CellMarker 2.0^144^ and Reactome^48^ databases.

#### Multiplex cytokine analysis

Supernatants from homogenised nasopharyngeal tissue were used to quantify cytokines and other soluble factors. IL-1α, IL-1β, IL-2, IL-3, IL-4, IL-5, IL-6, IL-9, IL-10, IL-12p40, IL-12p70, IL-13, IL-17A, CCL11, CSF2, CSF3, IFN-γ, CXCL1, CCL2, CCL3, CCL4, CCL5 and TNF-α were measured using the Bio-Plex Pro Mouse Cytokine 23-Plex Assay (Bio-Rad) according to the manufacturer’s instructions. Data were acquired on the Bio-Plex 200 system using BioPlex Manager™ 6.1 Software (Bio-Rad). Standard curve fitting and data analysis were conducted using Belysa Immunoassay Curve Fitting Software (Merck). CXCL2 and CXCL10 were measured using the mouse ELISA kit (Abcam), and IFN-α and IFN-β were quantified using the 2-plex ProcartaPlex™ Mouse IFN-α/IFN-β Panel (Thermo Fisher) as per the manufacturer’s instructions.

#### Immunohistochemistry analysis

Mouse heads were prepared and sectioned as previously described.^35^ In brief, 5-μm-thickness sections were prepared from formalin-fixed and decalcified mouse heads at rostral levels at approximately 200 μm.

Immunohistochemistry staining using a Leica Bond RX or DAKO Omnis auto stainer, whole slide scanning and image analysis/cell counting were conducted by the WEHI Advanced Histotechnology Facility. All sections were stained for Ly6G, CD4 and CD8 markers, using a DAB chromogenic protocol with amplification. Cell counts were conducted blindly within a region of interest containing the mucosa and submucosa, with the surrounding hard tissue structures as the limits and the intranasal cavities excluded from the analysis. Sections were scanned with an Olympus SlideView VS200 scanner, under brightfield mode and with a 20x air objective. Microscopic images were viewed and analysed using QuPath v0.5.1.^145^

## Statistical analysis

Graphs were plotted and statistical analyses were performed in R version 4.3.1 or GraphPad Prism V9.1.1. Data were expressed as medians ± IQRs, and groups were compared by the Mann-Whitney U test. The gene expression differential analysis between groups is tested using the moderated t-statistics and adjusted by Benjamini-Hochberg adjustment for multiple comparison as part of limma/voom in R.^139^ The p value of over-representation analysis of pathways or cell types was calculated by hypergeometric distribution with Benjamini-Hochberg adjustment for multiple comparison as part of ClusterProfiler in R.^140^

## Acknowledgments

We thank Susie Germano for support with the Bio-Rad BioPlex assay and Belysa Immunoassay Curve Fitting Software. We acknowledge the staff of the MCRI animal facility for their assistance with animal work. We thank the Advanced Histotechnology Facility at Walter and Eliza Hall Medical Institute for the resources, scientific contribution and technical expertise provided and Victorian Clinical Genetics Services (VCGS) for RNA sequencing. H.H. was supported by the China Scholarship Council - University of Melbourne PhD Scholarship. This study was supported by a Jack Brockhoff Foundation Early Career Medical Research grant (4212), a National Health and Medical Research Council (NHMRC) Ideas Grant (GNT1182442) and the Victorian Government’s Operational Infrastructure Support Program.

## Author contributions

Conceptualization: H.H., S.M., C.S. and O.W. Methodology: H.H., S.M., C.S., O.W. and A.M. Formal analysis: H.H., S.M., C.S., O.W. and A.M. Investigation: H.H., S.M. and A.M. Data curation: H.H. and S.M. Supervision: S.M., C.S. and O.W. Visualisation: H.H. and S.M. Resources: S.M., C.S., O.W., J.M. and S.P. Funding acquisition: S.M. and C.S. Writing-Original Draft Preparation: H.H. and S.M. Writing-Review & Editing: H.H., S.M., C.S., O.W., J.M., K.M., S.P. and A.M.

## Declaration of interests

SM and CS have received honoraria from Pfizer and MSD for presentations at symposia or attendance at expert advisory meetings unrelated to this study.

See figure document

## Supplementary Figures

**Figure S1:**
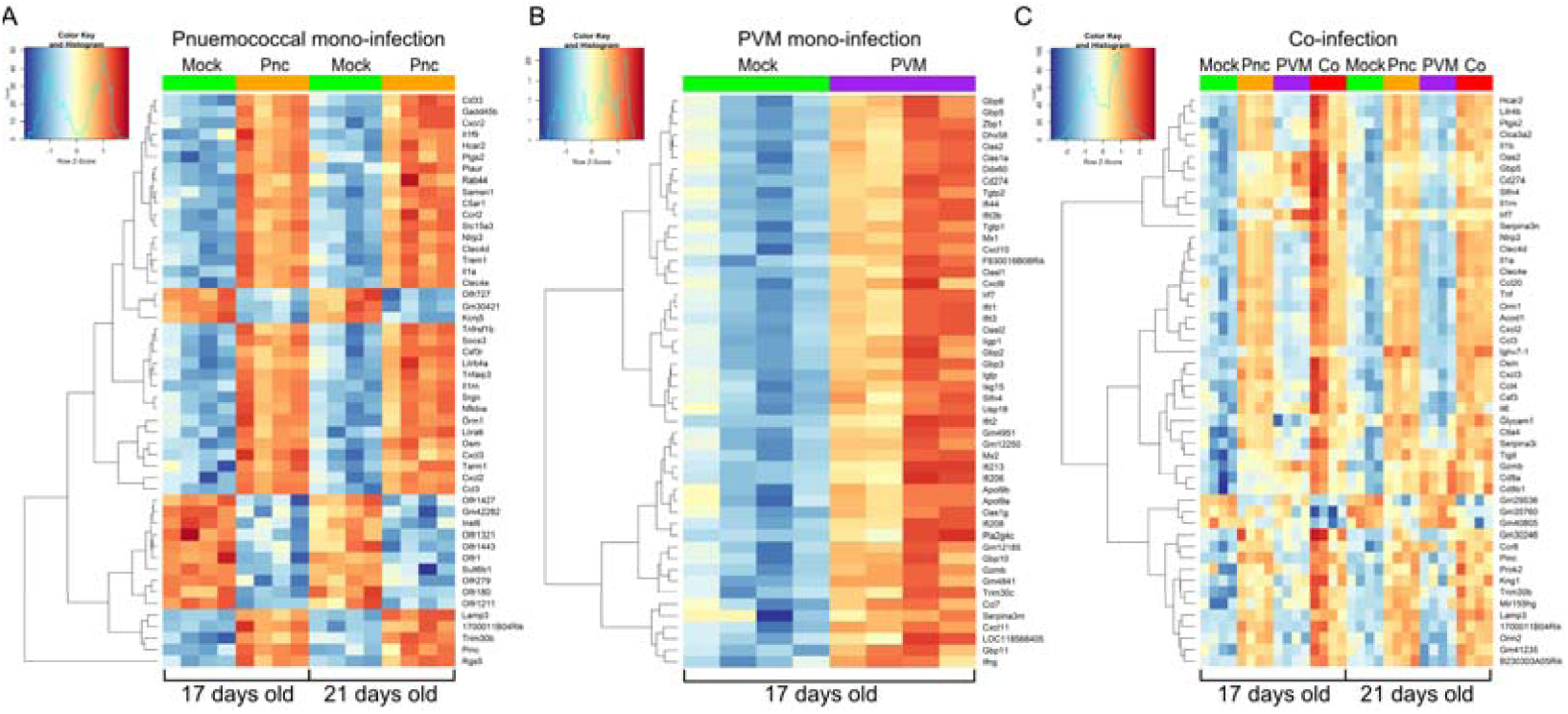
Heatmap of top 50 most differentially expressed genes by fold changes of pneumococcal mono-infection (A), PVM mono-infection (B) and co-infection (C). Colour key and histogram of Z scores were showed on the top-left corner. Dendrogram was plotted on the left to show the clustering of DEGs based on their expression patterns.

**Figure S2.**
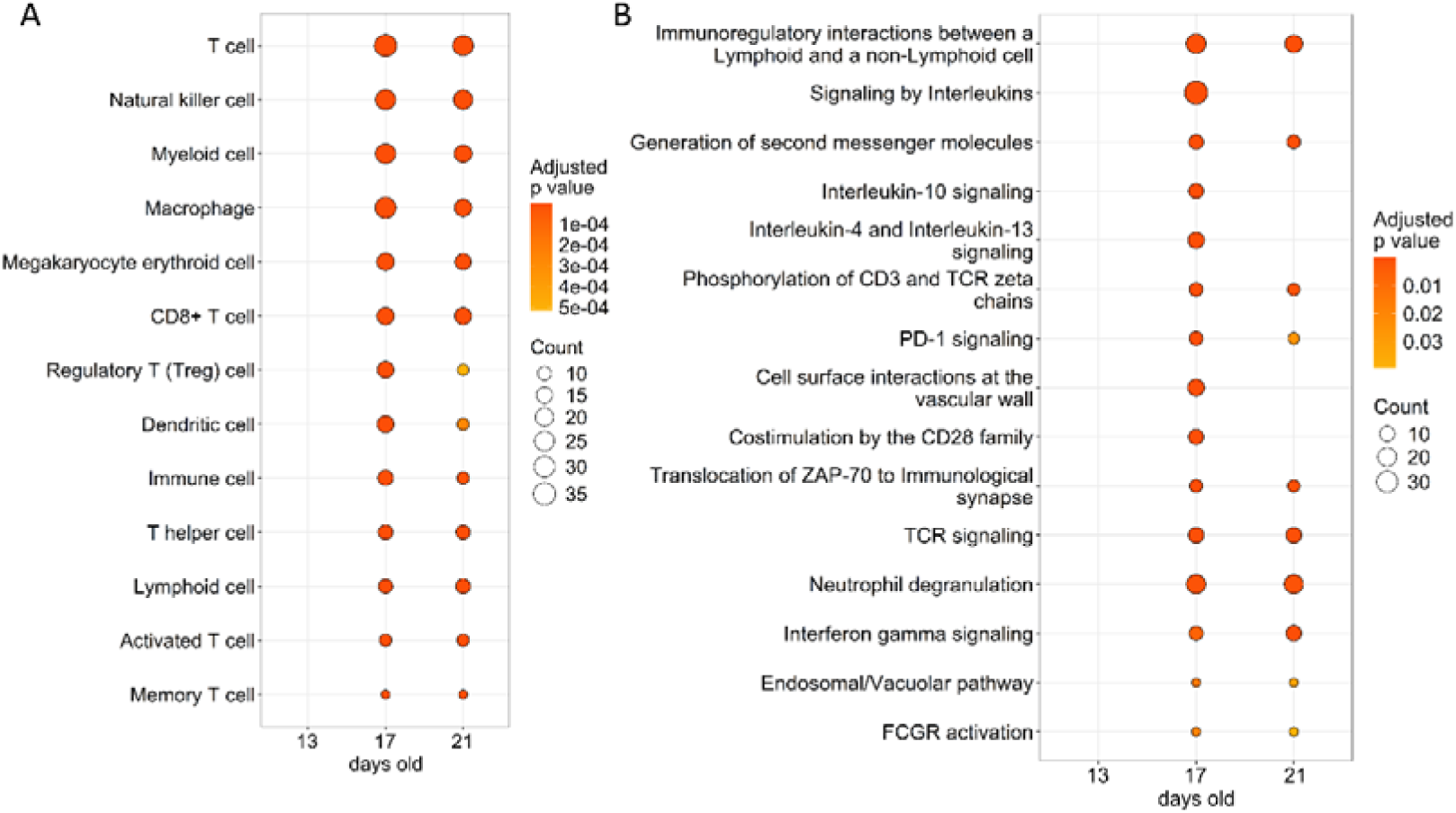
Over-representation analysis of unique DEGs during **co-infection for top enriched cell types (A) and pathways (B) at 13, 17 and 21 days old.** Over-representation analysis of DEGs uniquely found during co-infection for most enriched cell types (A) and pathways (B) at 13, 17 and 21 days old compared with mock-infected mice. The count indicates the number of DEGs identified within the gene set corresponding to each cell type or pathway in the reference database. Unique DEGs referred to genes that were only differentially expressed during co-infections but not mono-infections.

**Figure S3.**
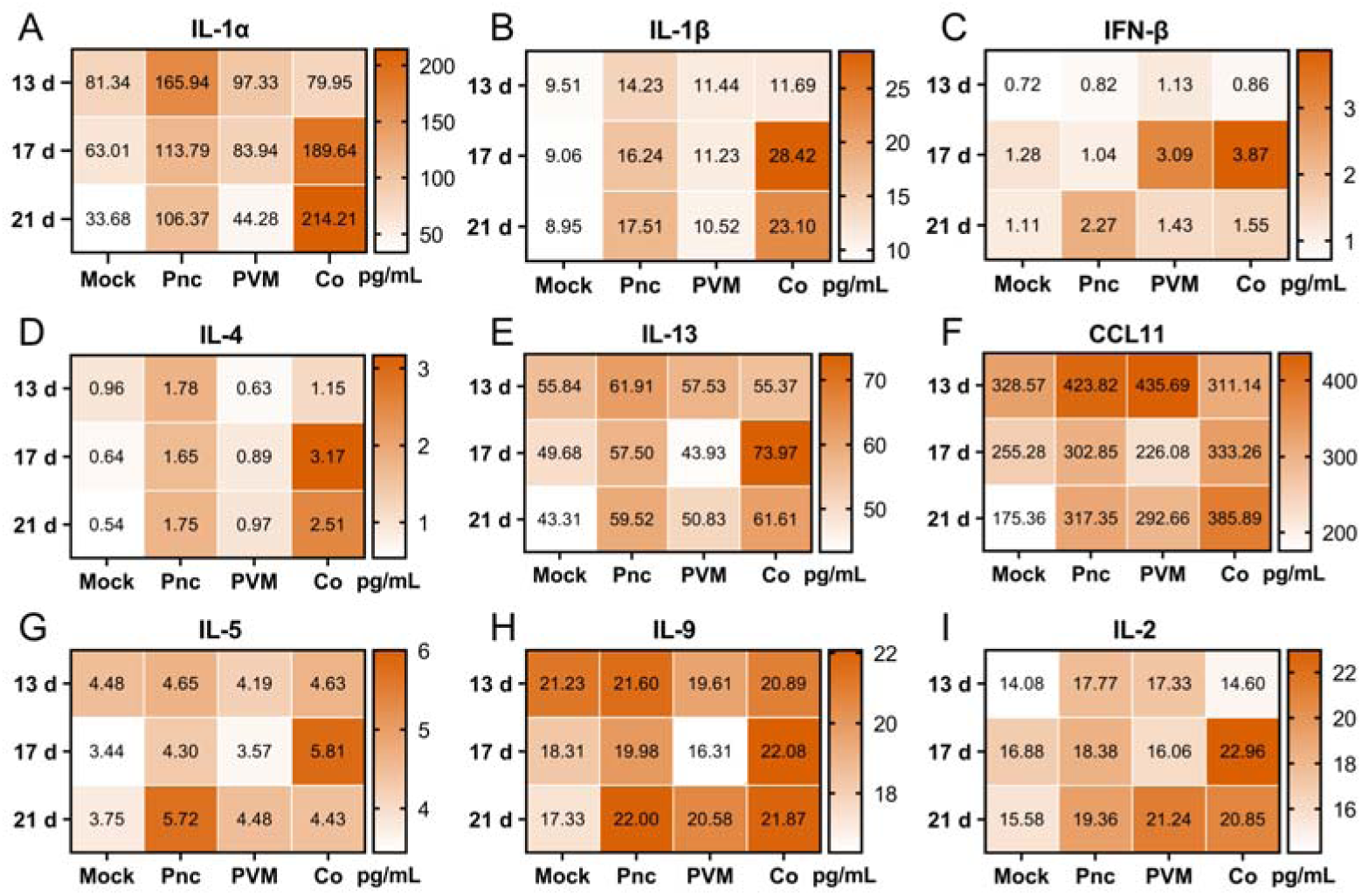
Levels of additional cytokines and chemokines during pneumococcal-PVM co-infection at over time. (A to I) Concentrations of critical cytokines and chemokines involved in mono-infections and co-infections at each time point. Concentrations of cytokines and chemokines in nasopharyngeal homogenates of 13, 17 or 21-day-old mice that were mock infected (Mock), pneumococcal mono-infected (Pnc), PVM mono-infected (PVM), or co-infected (Co). The colour of each block corresponds to the colour bar and represents the mean concentration of six samples per condition per time point.

## Supplementary tables

Table S1: DEGs during pneumococcal mono-infection

Table S2: DEGs during PVM mono-infection

Table S3: DEGs during co-infection

